# Life Under Pressure: Dissection of Cross-Phyla Metazoan Responses to Extreme Hydrostatic Pressure Reveals Pressure-Protective Heat Shock Acclimation

**DOI:** 10.64898/2026.07.06.736787

**Authors:** Mark E. Corkins, Anvita Bhattad, Tianyuan Hao, Nathaniel Williams, Mitchell P. Ford, Sean P. Colin, John H. Costello, Lance A. Davidson

## Abstract

The deepest ocean is one of the most extreme environments for life on our planet, combining near-freezing temperatures, low oxygen levels, and hydrostatic pressures reaching 111 MPa (1100 atm). Extreme pressures are predicted to alter many aspects of biology, including the physical properties of biological hydrogels, protein structure, and the solubility of gases in water. How organisms have adapted to live in these conditions is poorly understood. Studying these organisms *in situ* is difficult and requires specialized deep-sea equipment capable of withstanding the extreme pressure; raising these organisms in captivity is also challenging due to their extreme habitat requirements. Given these difficulties in studying deep-sea organisms, we set out to identify the problems shallow-dwelling organisms face due to increased pressure. These can provide insights into how organisms tolerate life in the deepest parts of the ocean. This project aims to take embryos of the shallow-dwelling aquatic organism *Xenopus laevis*, determine how surface-dwelling organisms fail under high hydrostatic pressure, and identify a means to survive this deadly pressure. We have designed a system to expose different embryonic stages of *X. laevis* to high pressures and observe its effects. After identifying the limits of survivability, we sought to understand how these embryos can acclimate to changing pressures. Comparative RNA-seq and cross-species analyses revealed a conserved, pressure-induced transcriptional response across phyla, with the heat shock pathway among the most strongly activated. Pre-activation of this pathway via prior pressure or other stressors enhances survival under otherwise lethal hydrostatic conditions.

## Introduction

The Ocean is stratified into 5 zones based on geography, light, temperature, and pressure. Each is populated by its own unique set of species that have evolved to live there. In the three deepest zones— midnight (Bathypelagic), Abyssal (abyssopelagic), and Hadal (trenches)—the temperature is consistently just above freezing (1). A major distinction among these three zones is hydrostatic pressure. Different species are restricted to specific zones (2) suggesting that pressure is an evolutionary constraint to which species adapt. For every kilometer descended in the ocean, the pressure increases by approximately 10 MPa (101.3 bar, 100 atm, or 1485 psi). The deepest part of the ocean reaches around 11 Km with a high hydrostatic pressure of 111 MPa (3). If removed from their adapted pressures, organisms suffer lethal physiological problems (4). *In vitro*, specific molecular changes have been observed or are predicted to occur due to changes in hydrostatic pressure, including membrane deformation (5, 6), alterations to hydrogen bonding that affect the way water interacts with DNA, RNA, and proteins, and changes in the solubility of gases (7-10). Very little is known about how these affect other aspects of biology or how shallow-dwelling organisms suffer from hydrostatic pressure.

Several studies have sought to identify the factors that have evolved to aid deep-sea organisms (11, 12). Most of these studies focus on genomic-level differences by comparing related species that are adapted to different depths. The best-studied of these are the snailfish of the family Liparidae, with many species that inhabit different ocean zones. Several snailfish species have had their genomes sequenced and partially annotated (13, 14). Comparative genomics suggests adaptation to high hydrostatic pressure includes increased osmolytes such as TMAO, altered lipid metabolism, and altered bone development (15). Currently, these observations are made from analysis of genomes, metabolic profiling, or physiological observations. None of these hypothetical adaptations has been tested *in vivo* to be pressure-protective.

Several studies of multicellular invertebrates have examined transcriptional changes due to changes in pressure (16-19). These studies are performed on less well-characterized organisms such as aquaculture-grown *Apostichopus japonicus* (sea cucumber) and wild-caught *Bathyacmaea lactea* (limpet) (16, 17, 20) looking at transcriptional changes of organisms at depths of around ∼3 km or on the surface. Given that these experiments were carried out in the ocean, differences between the deep-sea samples and those grown at sea level may be confounded by factors other than pressure, such as the food they eat, the composition of the water they live in, or the amount of light they are exposed to. Additionally, the invertebrates used in these studies have poorly annotated genomes, making whole-genome analysis difficult.

Humans, like all terrestrial organisms, have evolved to live at pressures found on the surface of the Earth from around 60 to 106 kPa, and develop medical complications when exposed to higher pressures (21, 22). Most complications arise because gases are more soluble in liquids under pressure, and when the pressure is reduced, dissolved gases effervesce to bubbles in various parts of the body. However, a range of clinical complications arises without a clear cause. For instance, high pressure can induce convulsions, presumably due to neurological dysfunction (23). Divers who spend too much time under high pressure can develop a related syndrome called high-pressure neurological syndrome (HPNS) (24). This syndrome is characterized by a variety of neurological conditions, such as headaches, tremors, and memory problems. These symptoms can emerge at depths as shallow as 100 m (1 MPa), far from the deep ocean floor at 10 km depth (25, 26). These clinical cases are likely due to multiple factors, including effects caused by increased hydrostatic pressure.

In this study, we utilize well-characterized, shallow-dwelling model organisms to investigate how animals acclimate to high hydrostatic pressure. We have assembled environmental chambers that allow us to culture aquatic organisms at specific temperatures and pressures in a controlled manner. We primarily use embryos of *Xenopus laevis* (African clawed frog) as a model of a shallow-dwelling amphibian. As there are no extant amphibians that can survive in the ocean, their evolutionary adaptation to the deep sea, if any, is likely limited. Additionally, *X. laevis* embryos are well-suited to study transcriptional changes, as they have a well-sequenced and annotated genome. *X. laevis* embryos are fully aquatic and do not contain air pockets, such as lungs or swim bladders. Early development is rapid, as they can progress from fertilization through neurulation in approximately 24 hours and form adult-like body plans by 4 days at room temperature, allowing for quick assessment of developmental progression and physiological function. These features allow us to pressurize embryos without concern that air-filled structures will rupture or force air bubbles into tissues upon decompression.

We began our study by exposing animals to high hydrostatic pressure and performing differential gene expression analysis with RNAseq. By comparing our dataset with those of other invertebrates, we found that the heat shock pathway is induced by pressures of 10 to 45 MPa across multiple species. We found that prior activation of the heat shock pathway by chemical and temperature shock increases an embryo’s tolerance to high hydrostatic pressure. Additionally, we found that suppression of this heat shock response (HSR) by injection of a competitive inhibitor for a key transcription factor (Hsf1) decreases tolerance for hydrostatic pressure. This study demonstrates that HSR is a key pathway that underlies metazoan acclimation to extreme hydrostatic pressure.

## Results

### Tolerance to Pressure

To test how *X. laevis* responds to hydrostatic pressure, we first sought to determine the range of conditions that embryos could survive. To accomplish this, we engineered a pressure environmental chamber to culture embryos in pressures up to 140 MPa. This system consists of a programmable syringe connected to an environmental chamber constructed from 316SS tubing (Fig. S1 and Methods). For some experiments, we isolated embryos in compartments composed of hydrated PVC tubing with PDMS plugs. We did not observe any differences in pressure survival between embryos cultured in the chamber alone or in tubing. These experiments were performed on blastula-staged embryos (NF st. 8-9) (27, 28) as they are older than the maternal-to-zygotic transition (MZT), allowing the embryos to acclimate transcriptionally (29). The choice of a post-MZT stage was important because it allowed us to measure changes in gene expression using methods such as RT-PCR or deep sequencing. Additionally, embryos at this stage are just beginning to gastrulate, a period during which the embryo undergoes rapid cellular movements, allowing us to synchronize their developmental stage (30).

After preliminary experiments, we settled on a standard protocol of a 30-minute gradual ramp to the indicated pressure (Ramp phase), hold at the indicated pressure for 2 hours (Hold phase), then return to lab pressure over 15 minutes (Recovery phase) (Fig. 1A). The 30-minute ramp phase was chosen as it is slow enough to not cause barotrauma, physical tissue damage caused by rapid pressure changes, but is suffiently fast to limit transcriptional responses to high pressure. The 2-hour hold phase was long enough for pressure to cause damage, but not so long that secondary effects, such as hypoxia, could affect viability. A 15-minute recovery phase was chosen because, when we tested whether the duration of the recovery phase affects survivability, we did not see a strong effect (Fig 1C-E). Therefore, we reduced the pressure as quickly as the pump could do so accurately, without causing barotrauma. To test for viability, recovered embryos were cultured at 14°C overnight. At maximal pressures below 35 MPa, embryos survived with no noticeable developmental defects (Fig. 1B). However, at pressures above 40 MPa, embryos appeared normal upon removal from the system but died of cell lysis within a few hours (Fig Sup Movie 1). In some clutches of embryos, pressure increased the frequency of exogastrulation. The underlying causes of variable exogastrulation are currently unclear; many stressors, such as osmotic stress, can induce exogastrulation, especially when egg quality is poor (31). Within a single exposure to pressure, we observed that either >95% died or >95% survived and developed relatively normally. We could not find conditions under which approximately half of the batch survived. Additionally, over the summer months, pressure tolerance increased, with some batches surviving above 60 MPa, suggesting that the maternal environment affects the pressure resistance of their offspring. Although our adult animals are housed in a temperature- and light-controlled room and reared in RO water with added salt, seasonal variation in egg quality is a common observation in the *Xenopus* community, suggesting that animals can still sense external seasonality. Due to this variability, all experiments were performed in triplicate and, when possible, with matched controls from the same clutch of embryos. Additionally, to prevent confounding temperature effects, all embryos were raised at 14°C and kept at 14°C when not being experimentally manipulated.

**Figure 1:**
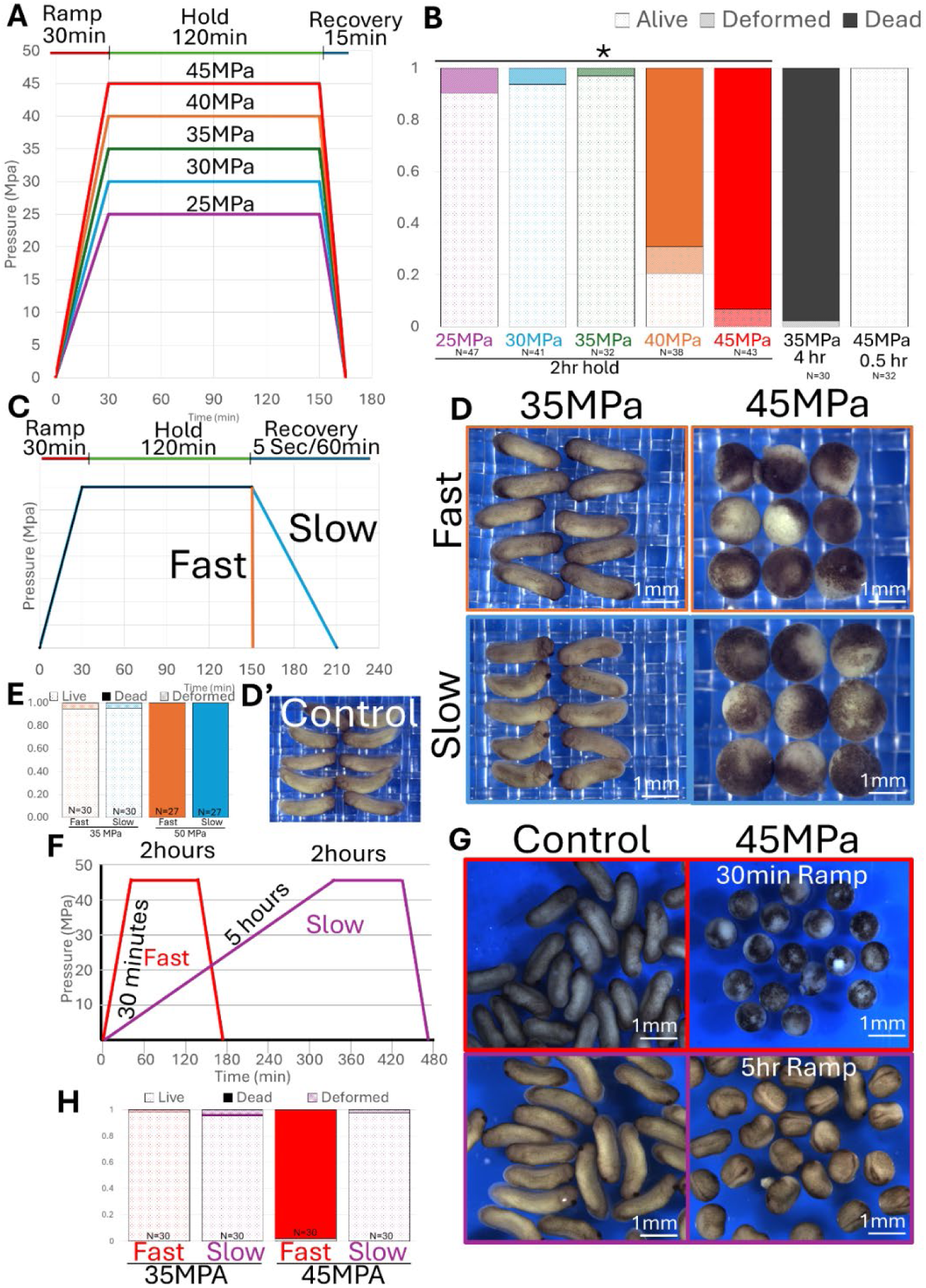
*X. laevis* is sensitive to high levels of pressure. **A,B)** The level of pressure and the amount of time held at pressure are important for survivability. Embryos were treated according to panel A. The graph indicates the fraction of embryos that survive each condition. **C-E)** The rate of recovery (depressurization) does not affect survivability. **D)** Images of embryos the day after embryos were exposed to pressure, as diagrammed in C. **F-H)**Time to pressure is important for survivability. **G)** Images of embryos the day after embryos were exposed to pressure, as diagrammed in C. **A, C, F)** Diagram showing experimental conditions to test for lethality. Embryos were grown to stages 8-10 and treated with pressure as shown in the diagram. **B, E, H)** Graphs indicating the percentage of embryos surviving treatment diagrammed by A, C, and F, respectively. * p<0.05 by Fisher’s Exact test

We also experimentally tested the effects of each phase of the pressure cycle, including the duration of the hold phase. Embryos that were kept at 25 MPa for 5 hours resulted in embryo death (Fig. 1B). However, when we lowered the pressure to 10 MPa, we could maintain pressure for up to 12 hours with minimal long-term harm. However, 10 MPa for 20 hours resulted in lethality. Additionally, higher pressure for shorter durations, such as 45 MPa for 1 hour, was survivable. This indicates that both the duration and the magnitude of pressure contribute to lethality.

As a final parameter, we sought to determine whether ramp time (the time to reach maximum pressure) affects survivability. We exposed embryos to 45 MPa of pressure as described above, and instead of reaching 45 MPa in 30 minutes, we extended the ramp to 5 hours (Fig 1F). This longer ramp allowed embryos to survive this treatment (Fig 1 GH). Overall, our data indicate that the time to reach pressure, the pressure level, and the duration at pressure have strong effects on survival, whereas recovery time does not. Additionally, survival of embryos exposed to high pressure after a 5-hour ramp suggests that embryos can acclimate to pressure.

### Development

Extended culture of *X. laevis* embryos in our environmental chambers can result in a developmental delay (Fig 1G). Although many stressors, such as heat shock, can induce short developmental delays of up to a few hours (32), we observed delays lasting up to 20 hours. The primary cause appears to be the steel endcap and cylindrical tube chambers, since embryos exposed to 316SS and tubing exhibit a pressure-independent delay (Fig S2A). Confoundingly, embryos cultured in the individual parts of the stainless steel chamber do not exhibit developmental delay (Fig S2B,C). The developmental delay does not affect our later results, as embryos cultured in chambers for 5 hours do not induce a heat shock response (Fig. S2D). Therefore, we do not predict that this delay will affect our findings in relation to pressure, but note that embryos cultured conventionally were at later stages of development.

Since the culture space in the sealed, pressurized environmental chamber is a long cylindrical pipe and the entire system is limited to ∼70 ml, we suspected that hypoxia might contribute to the developmental delay. To test this possibility, we conducted a series of experiments in which we varied the amount of air to which embryos were exposed. We conducted a pressure experiment using our pump to introduce excess air into the fluid. Two sets of embryos were placed in the pressure chamber. One set was open to the system and exposed to hyperoxic conditions, while the other set was sealed in flexible PVDF tubing, as shown in Figure S1C, thereby separating the embryos from the environment. Both sets showed a similar developmental delay (Fig S3B). We also removed oxygen from the media by either bubbling nitrogen or degassing under a vacuum. These embryos developed at a rate similar to untreated embryos, suggesting that dissolved gas levels do not strongly affect developmental rate in early embryos.

### Cytoskeleton

Many components of the cytoskeleton are reportedly sensitive to high pressure, with microtubules among the most sensitive (33-36). Given the cytoskeleton’s role in many cellular processes, we sought to examine its potential contribution to lytic death. We prepared embryos by injecting reporters for the primary cytoskeletal components emtb::mCherry (microtubules), lifeact::miRFP670 (F-actin), and krt8::EGFP (keratin intermediate filaments). Embryos were subjected to 35 MPa of pressure (37-39). After the 15 minute recovery, they were immediately fixed with 3.7% PFA in MBS for 5 minutes and imaged. Although ectoderm cell shapes were more rounded, we did not observe major disruptions to the microtubule or F-actin networks; however, the keratin network was strongly disrupted (Fig S4). This was confirmed by antibody staining. Keratin knockdown in many species leads to developmental defects, some of which are lethal; however, at the cellular level, keratin is not considered necessary for survival (40, 41).

### Transcriptional response

Following these candidate approaches, we next sought an unbiased approach to identify how embryos respond to pressure and possibly understand why they die. Considering that, at the blastula stage, embryos would survive if slowly brought to 45 MPa but would not under a 15 minute ramp, suggests that embryos could acclimate. Survival after a 5 hour ramp duration suggests that acclimation is slow and involves transcriptional induction. To identify genes regulated by pressure, we performed RNA-seq analysis of pressurized embryos. Given the developmental delay, two controls were used. One set of embryos was grown at 14°C to delay development to more closely match the pressurized embryos, while another set was grown at room temperature, matching the temperature of the pressurized embryos. Embryos were brought up to 45 MPa over 5 hours, held at 45 MPa for two hours, then returned to normal atmospheric pressure over 15 minutes (Fig 2A). Embryos were frozen at -80°C, and total RNA was extracted from all embryos at the same time.

**Figure 2:**
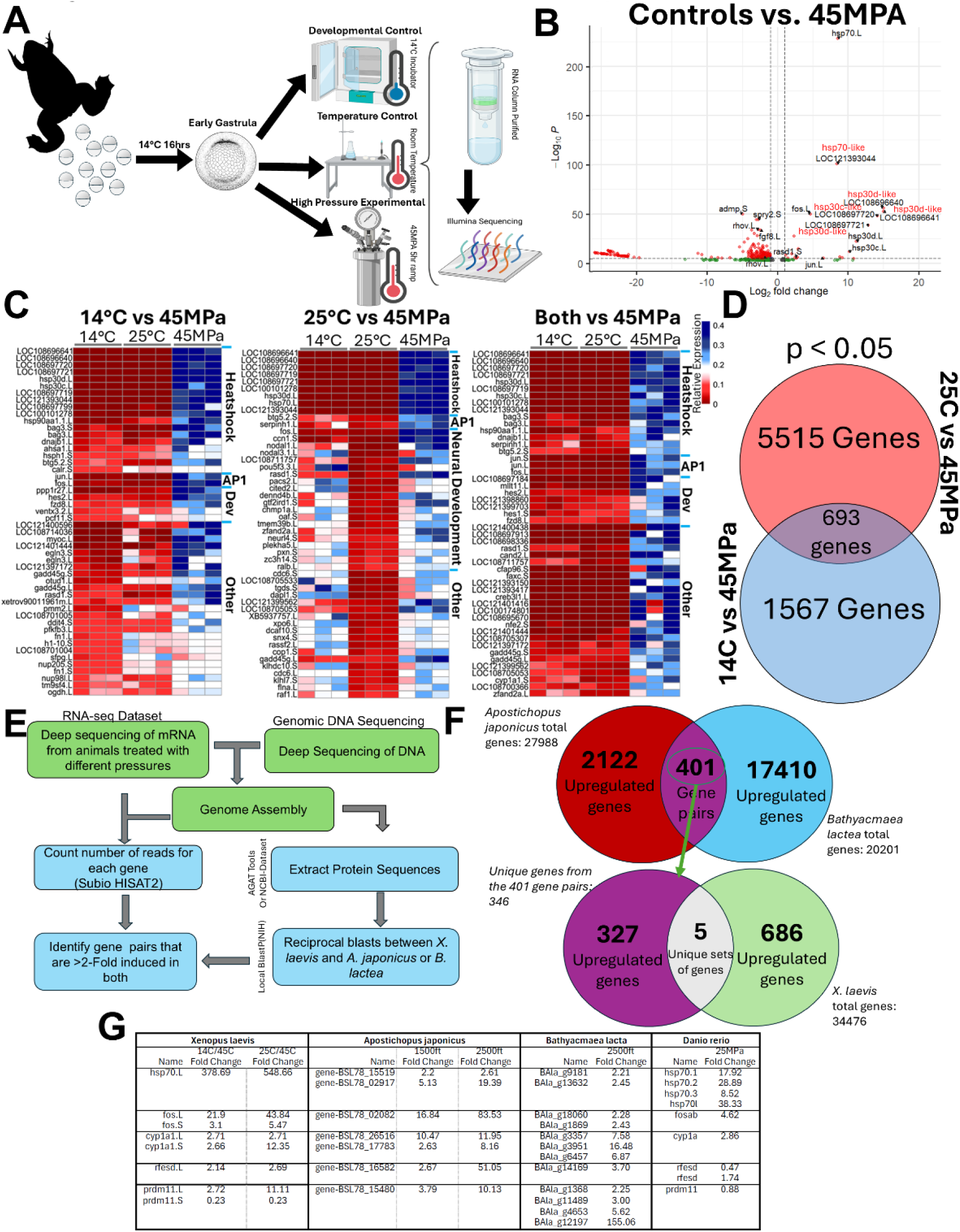
RNAseq experiments identify gene expression conserved across metazoans. **A)** Diagram representing experimental conditions used for *X. laevis* embryos**. B)** Volcano plot illustrating genes induced by pressure in *X. laevis*. The gene family for genes with no assigned gene name is indicated in red. **C)** Heatmap of the top 50 most significantly induced genes. Genes are sorted by category, listed on the right of the figure, and by fold change. The label above indicates which control is used. **D)** Venn diagram illustrating the amount of overlap between the use of the temperature control, and developmental control datasets. **E)** Diagram of the workflow to find universally upregulated genes. Green steps for *A. japonicus* and *B lactea* were done by (16, 17). **F)** Venn diagram of positive gene pairs identified. Overlap indicates gene pairs in which both genes are at least 2-fold induced. **G)** Table showing fold induction of positive hits across species.

Pressure can inhibit transcription (42) and may not affect all transcripts similarly; also, small differences in developmental timing shortly after the maternal-to-zygotic transition (MZT) could affect the gene expression profile. To try and account for this possibility, we normalized the raw counts using multiple methods. The first data set was normalized using the standard technique of calculating Transcripts Per Million (TPM). Secondly, to account for the effects of pressure on developmentally regulated transcription, data were also normalized to a subset of 8 genes that were found not to change during gastrulation, including *clta.L*, *lpcat3.S*, *slc35b1.L*, *mtch2.L*, *mcts1.L*, *ralb.S*, *cox7b.S*, *and prcp.S* (43, 44). The total counts of these genes were summed up for each sample, and the counts were normalized to the 8-gene subset. Both normalization methods yielded similar results and did not alter the top 50 results. 693 genes were found to have p-values <0.05 in both the 14°C control and the room-temperature control datasets (Supplemental Table 1). The largest set of genes induced by pressure comprises a family of molecular chaperones associated with the heat shock response pathway and a set of genes expressed in developing neural tissues (Fig 2B,C).

To identify genes universally upregulated by pressure in metazoans, we started by reanalyzing previously published RNA-seq datasets from invertebrates housed at sea level or caged at greater ocean depths. One dataset from the shallow-water sea cucumber *Apostichopus japonicus* (15 and 25 MPa), and a second from the deep-sea limpet *Bathyacmaea lactea* (∼14.2 MPa) (16, 17). The raw FASTQ files were reprocessed using the same pipeline used for the *X. laevis* RNA-seq experiment. Given that neither invertebrate organism had a well-annotated genome, homologous gene pairs were identified using reciprocal BLAST (Fig 2E). We found that 401 gene pairs, comprising 346 unique genes, were upregulated by at least 2-fold in both species (Fig. 2F). To further reduce this number, the genes identified in invertebrates were used in a reciprocal BLAST search against *X. laevis* to identify universally induced pathways. Only 5 genes were upregulated in all three species (*cyp1a1*, *prdm11*, *hsp70*, *rfesd*, and *fos;* Fig 2G). To further validate these as universally upregulated across a wide range of species, we pressurized both Zebrafish (*Danio rerio*) and Moon Jellyfish (*Aurelia aurita*) to 25MPa for 2 h, isolated RNA, and carried out RNA-seq (Supplemental Table 1). In Zebrafish, 4 copies of *hsp70*, *fosab*, and *cyp1a* were upregulated in response to pressure; however, *rfesd* and *prdm11* were not. Because the Moon Jellyfish genome is poorly annotated, many genes are missing; however, *hsp70* was strongly induced. This leads us to conclude that hsp70, cyp1a, and fos are induced in response to pressure in a wide range of organisms across the animal kingdom.

As *hsp70* and fos are considered direct targets of the heat shock response pathway (HSR) (45), and though not direct targets, cytochrome P450 proteins such as Cyp1a1 and Rfesd are frequently associated with responses to stressors, including heat shock (46-48). Moreover, the most induced genes in *X. laevis* were eight variants of the *hsp30* gene, *hsp70*, and *hsp90aa1,* all members of the HSR pathway (49-51). Therefore, we focused on the HSR pathway to determine whether it mediates tolerance to high pressure.

### Heatshock regulates pressure resistance

To determine whether pressure resistance involved HSR, we adopted strategies to induce a heat shock response independent of pressure and then assessed whether this pretreatment could enable embryos to survive normally lethal pressure levels. We activated HSR using the well-characterized heat-stress induction technique (52). Heat shock was performed in 60 mm dishes on a temperature-controlled hot plate. Embryos were exposed to 32°C for 1 hour, then to 34°C for 1 hour, and finally to 37°C for 30 minutes. Embryos were then exposed to 45 MPa of pressure, which is typically lethal. The embryos that were heat-shocked survived, whereas control embryos did not (Fig 3A). Exposure of embryos to heat and pressure leads to developmental abnormalities; this was expected, as heat shock is known to cause developmental abnormalities (53). Therefore, we also activated HSR using the heavy metal cadmium, which is reported to activate the HSR pathway in *Xenopus* (54, 55). Similar to heat stress, treatment for 2.5 hrs with 50 µM CdCl_2_ was also pressure-protective but did not generate developmental abnormalities (Fig 3A). Without pretreatment, exposure of embryos to heat and pressure concurrently reduced survivability (Fig 3B). Thus, transcriptional activation of the heat shock response pathway underlies the tolerance to high pressure seen in embryos subjected to a slow ramp.

**Figure 3:**
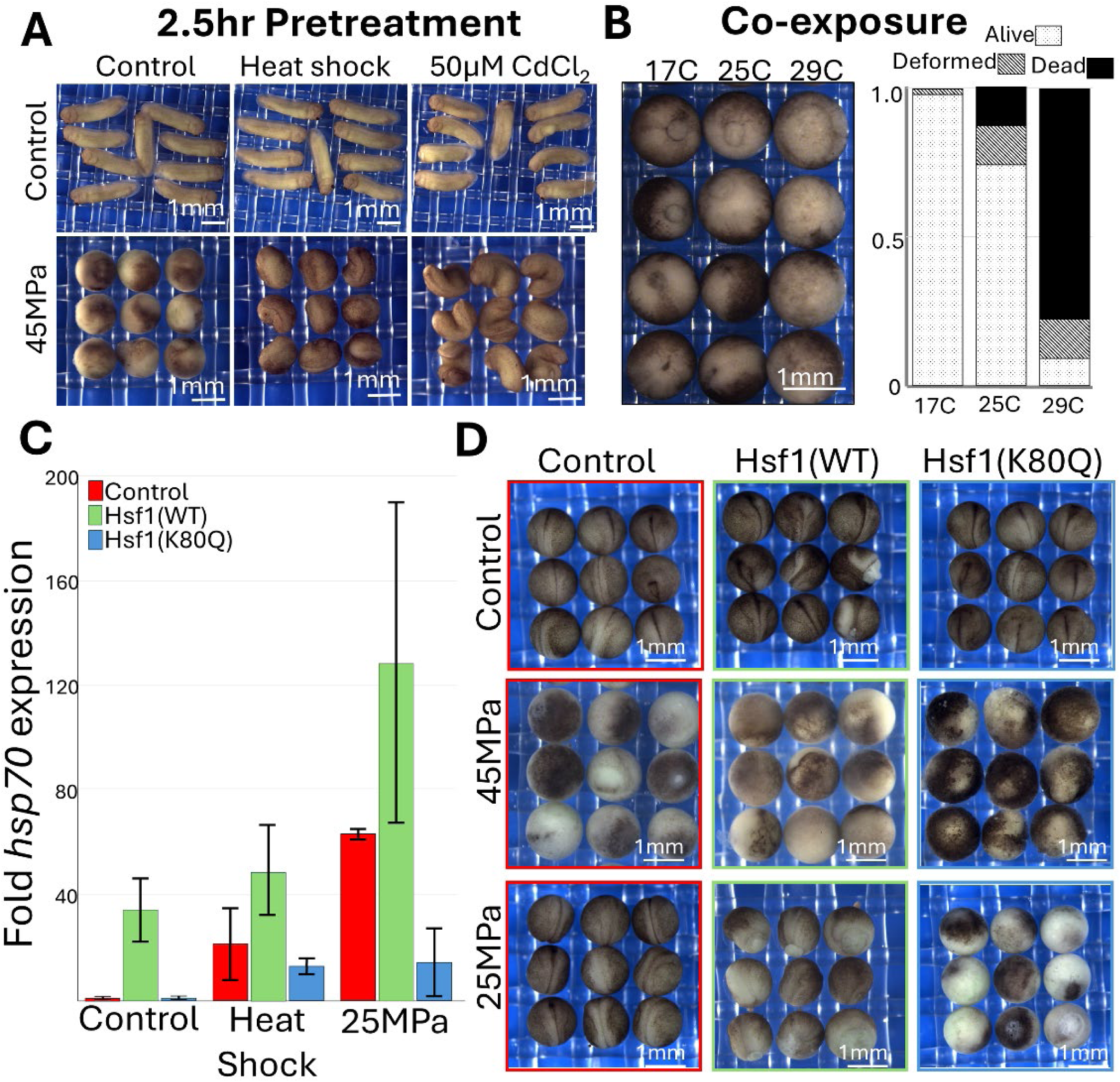
HSR is important for surviving hydrostatic pressure. **A)** Embryos were pretreated with either heat stress or cadmium stress, then treated with 45 MPa of pressure. Embryos were grown at 14°C, then imaged the next day. **B)** Embryos were exposed to 20 MPa of pressure for 5 hours at various temperatures, then imaged after pressure returned to atmospheric pressure. **C)** Embryos were injected with mRNA encoding either xlHsf1 or xlHsf1(K80Q), then treated with either heat or pressure, and expression of *hsp70* was measured by real-time PCR. Results are shown as fold induction relative to control conditions. **D)** Embryos were injected with the indicated mRNA, then exposed to the indicated pressure. Embryos were allowed to grow overnight at 14°C, then imaged.

Since activation of HSR increased tolerance to pressure, we wondered if suppression of the HSR would decrease tolerance. To accomplish this, we expressed a competitive inhibitor of the core HSR transcription factor Hsf1. Human HSF1 was tested and found to be nonfunctional in *X. laevis*. Therefore, we cloned *X. laevis hsf1.L* and fused it to a fluorescent protein (Hsf1::EGFP). Overexpression of Hsf1 constitutively induced *hsp70* expression (Fig 3C). Before gastrulation, Hsf1::EGFP is primarily localized to the cytoplasm; after neurulation, Hsf1 is constitutively nuclear localized in the ectoderm (Fig S5A). Our observations are consistent with previous reports that HSR is suppressed prior to gastrulation (56). In mammals, acetylation of lysine-80 in Hsf1 is required for proper DNA binding. As Hsf1 functions as a homo-trimer, incorporation of Hsf1(K80Q) is able to form a trimer but is unable to bind to DNA, acting as a competitive inhibitor (57). Therefore, the mutant Hsf1(K80Q) functions as a competitive inhibitor of endogenous Hsf1 (58). We verified that this construct suppressed heat shock-induced transcription of hsp70 as well as suppressed the effect of overexpressed Hsf1(WT). (Fig 3C, and S5B).

To test if Hsf1 plays a role in pressure sensitivity, embryos were injected with either the wildtype or the competitive inhibitor Hsf1(K80Q). Expression of wildtype Hsf1 had no effect on survivability but did result in developmental defects, such as exogastrulation (Fig 3D). Embryos injected with Hsf1(K80Q) were sensitized to high pressure and died when exposed to normally survivable 25 MPa Indicating that activation of the heat shock pathway is necessary but not sufficient to increase pressure tolerance. This finding suggests HSR pathway works in parallel with other pathways to enable pressure tolerance.

### Tadpole muscle paralysis

Experiments up to this point were performed on early blastula-stage embryos, and we were curious about the effects on more developed, e.g., physiologically functioning organisms. Therefore, we examined the effects of high pressure on tadpole-stage embryos (st. 40, ∼3 dpf at 25°C) since these have functional nervous, muscle, circulatory, and renal systems. These embryos were more sensitive to pressure, with pressures above 30 MPa being lethal, whereas the blastula stages can survive 40 MPa. As in early-stage embryos, death occurs through gradual cell lysis, with the obvious sign being visible lysis of skin cells (Fig Sup Movie 2). At the survivable pressure of 25 MPa, the most prominent phenotype observed was paralysis. Paralyzed embryos have a visible dorsal curvature, their hearts are stopped, and they do not respond to touch. After approximately 30 minutes, the heart would restart, and by the next day, they had recovered completely with no apparent developmental or physiological abnormalities (Fig 4A). To investigate this paralysis phenotype further, we questioned whether the paralysis was due to muscle damage. To test this, tadpoles were stained with phalloidin to label F-actin. We focused on the somitic myotome because it is a well-organized skeletal muscle in early tadpole stages. Pressure-treated tadpoles retain individual sarcomeres, but muscle fibers appear thinner and less straight than in controls (Fig. 4B). Additionally, small actin-positive particles were also visible around the myotome after pressure.

**Figure 4:**
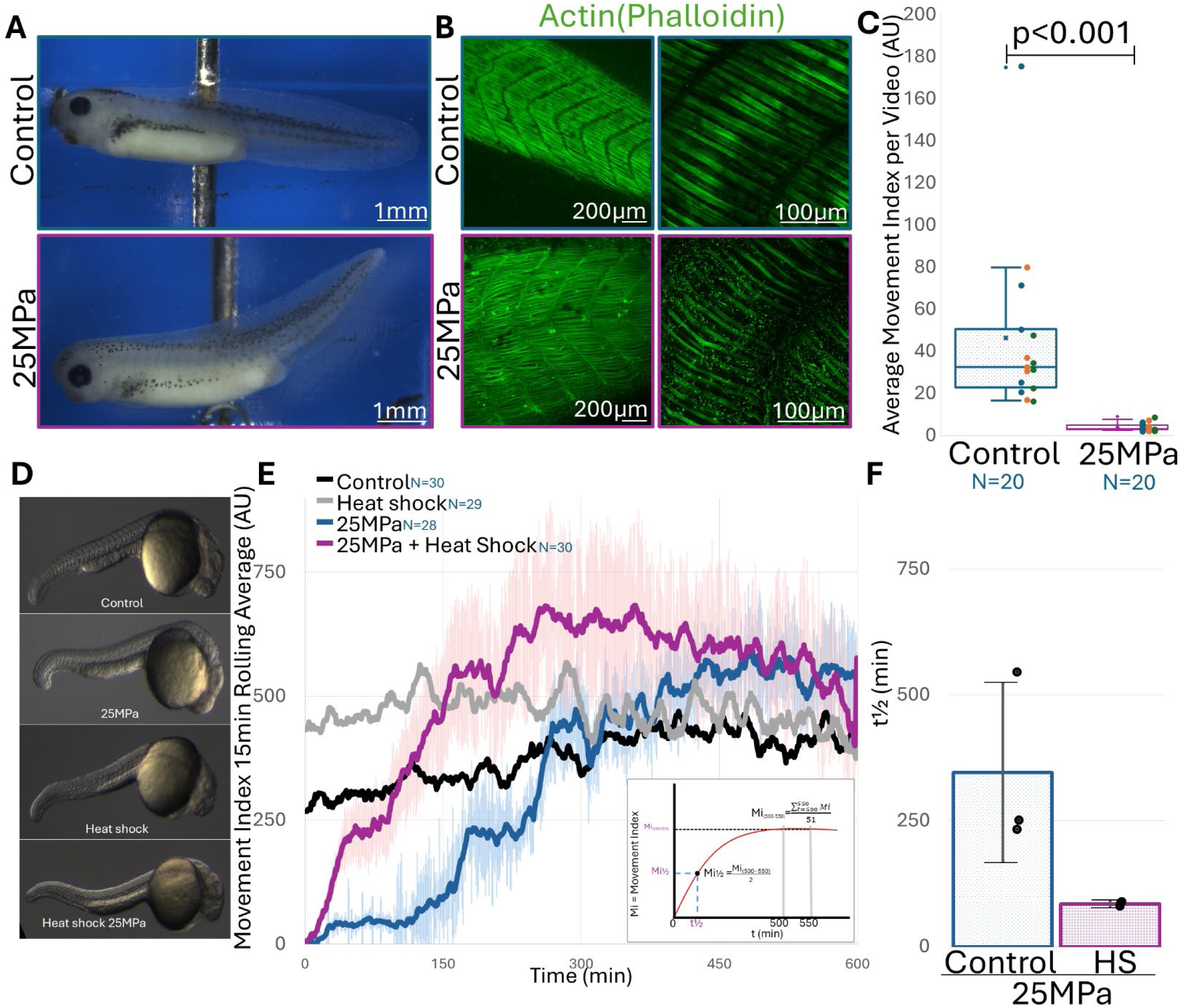
Pressure has a paralytic effect on tadpoles, which can be partially rescued by the heat shock pathway. **A)** Image of a tadpole shortly after being treated with either normal atmospheric pressure (Blue) or 25 MPa (Purple). **B)** Image of myotome stained with phalloidin after treatment with pressure. The right image is a more magnified view of the muscle. **C)** Embryos were electrically stimulated using a TENS device, and the movement of embryos was quantified. Box plot of the average intensity across multiple timelapses. The box plot shows the average movement index across all timelapses, while dots indicate the movement index for each individual timelapse. Different colored dots indicate different days and batches of embryos. **D)** Representative images of Zebrafish after exposure to 25MPa of pressure. **E)** Movement index of embryos after exposure to pressure. **F)** Bar chart showing the amount of time it takes to return to half mobility. Blue lines/boxes are embryos exposed to 25MPa of pressure, whereas purple indicates embryos heat-shocked prior to exposure to 25MPa.

While the myotome appeared damaged, we wanted a more direct test of functionality. We used a transcutaneous electrical nerve stimulation (TENS) device to stimulate muscle contraction by delivering electrical pulses. To prevent untreated animals from swimming away, they were anesthetized with 0.005% MS222 for 15 minutes prior to treatment. The animals were washed in anesthetic-free 1/3x MBS and placed over an electrode embedded in PDMS; they were subject to stimulation and recovered in under 10 minutes after removal from anesthesia. We did not anesthetize pressure-treated animals, as, due to paralysis, they did not swim away. We were also interested in whether we could observe any movement. Clusters of electrical pulses were delivered to allow the muscles time to relax, and in the movement analysis, we could observe that the movement was due to the muscle stimulation, rather than other motions of the system. Control animals would bend in response to stimulation, and this movement was coincident with the current pulses (Figure S7). Pressure-treated animals failed to respond to electrical stimuli, and no muscle contraction was visible in response to current. (Fig 4C). Movement was measured by an ImageJ algorithm described in Figure S6. The small amounts of movement we measured were largely due to beating cilia, which continued in both MS222 and pressure-treated tadpoles. Additionally, the continuing heartbeat and bubbles generated by the electrical current were minor contributors to the measured movement. We find that ciliary beat continues after exposure to high pressure, whereas the ability to contract muscles is lost upon exposure (Fig. S3 Movie 3).

*X. laevis* is largely stationary until around stage 43 (mid-tadpole stages). They will move if touched but remain still if they are undisturbed, making it hard to measure when they recover. Therefore, to investigate how long paralysis persists, we used Zebrafish (*Danio rerio*). Zebrafish are highly motile at 1dpf and, at these early stages, exhibit all of the same benefits described earlier for *X. laevis*. We observed that their pressure tolerances were similar to those of tadpole stage *X. laevis*, and they upregulated heat-shock genes in response to pressure (Fig 2G and Table S1). Using the same electrostimulation protocols applied to *X. laevis* tadpoles, we treated 1 dpf Zebrafish with 25MPa of pressure and observed a similar paralysis phenotype. Similar to *X. laevis,* there were no observable differences seen between pressurized and control animals, other than that pressurized animals were immobile (Fig 4D). Using time-lapse imaging, we applied the same movement macro as in the *X. laevis* electrostimulation experiments (Fig Sup Movie 4). We saw that movement gradually returned to control levels in approximately 6.5 hours (Fig 4E). We also noticed that movement in the control animals was, on average, relatively constant throughout the time-lapse.

Given that we found that heat shock pretreatment increased survival to the blastula stage in *X. laevis*, we asked whether heat pretreatment could allow animals to recover faster. To test this, we heat-shocked 1 dpf Zebrafish embryos and subjected them to 25 MPa of pressure. Then we asked whether the recovery time had been shortened. To measure recovery, we calculated the ½ time recovery (t_½_); see materials and methods for details on calculations. As the unpressurized embryos did not show a reduction in movement and therefore did not recover, we did not calculate it for those samples. However, both control and heat-shocked animals had similar levels of movement. For the pressurized embryos, the untreated samples have a t_½_ of 345 minutes, and for the heat-shocked embryos, that time drops to 84 minutes (Fig 4F). Indicating that, similar to what was seen for the blastula-staged embryos, pretreatment with heat shock significantly improves recovery from pressure.

## Discussion

Just like terrestrial environments, the ocean contains many biomes with different environmental conditions, including temperature, light, and pressure. Understanding how animals tolerate and adapt to environments is important for understanding how they cope with ever-changing conditions, such as those experienced by animals undergoing ocean acidification or diving deeper as the ocean warms (59-61). This work examines the effects of high hydrostatic pressure on whole metazoan organisms in a controlled lab environment while seeking to exclude extraneous variables such as temperature, water composition, and the effects of air under pressure. In this study, we sought to understand how vertebrates succumb, acclimate, and tolerate exposure to high hydrostatic pressure, such as that found in the deepest ocean. We focused our study on embryos of the near-surface-dwelling organism *X. laevis*. As a first step, we set out to identify which pressures are lethal. Then asked which part of this process was important to survivability: Ramp Hold and/or Recovery. We found that the duration of high-pressure exposure (hold time) and the time to reach pressure (ramp time) were critical variables in determining the point at which embryos tolerate the toxic effects of pressure. We found that pressure of around 40 MPa was the point at which *X. laevis* succumbed, and cell lysis occurred shortly after removal from the system. However, our findings show that a slow ramp allows embryos to acclimate to and survive normally lethal pressures.

After a transcriptional analysis, we found that 693 genes were regulated by pressure. To identify genes and pathways universally induced in metazoans under pressure, we used published deep sequencing datasets to compare genes across multiple species. We found 5 unique sets of genes were induced in all three organisms: the limpet *B. lactea*, the sea cucumber *A. japonicus*, and the frog *X. laevis* (*cyp1a*, *prdm11*, *hsp70*, *rfesd*, and *fos).* After the addition of Zebrafish and Moon Jellyfish RNA-seq datasets, this list was reduced to just (*cyp1a*, *hsp70*, and *fos).* This data, along with the observation that the core HSR genes hsp30, hsp70, and hsp90aa were strongly enriched in the *X. laevis* RNA-seq data, directed us to further study the HSR pathway.

We found HSR is a key pathway that allows embryos to acclimate to pressure. We both activated and suppressed the HSR pathway, finding increased and decreased tolerance to pressure, respectively. By pre-treating embryos with heat shock and heat-shock inducing CdCl_2_ treatment, embryos were able to tolerate normally lethal pressures with a short ramp time. Alternatively, by suppressing the HSR pathway with competitive inhibitors of the core transcription factor of the HSR pathway, we made embryos more sensitive to pressure, indicating that the HSR pathway is key to pressure tolerance. Though many studies have shown that *hsp70* is induced by hydrostatic pressure (62-64). Ours is the first to demonstrate that the HSR pathway has a role in pressure tolerance.

We took this a step further and examined how physiologically functional *X. laevis* tadpoles tolerate pressure. We observed that pressure transiently paralyzed tadpoles. This is likely due in part to muscle damage, as animals did not respond to direct neuromuscular stimulation using a TENS device. To measure recovery time, we used larval Zebrafish, as they are more mobile than *X. laevis* embryos but remain confined to the vitelline membrane. Not only did we find that it takes around 6.5 hrs to fully recover to control levels, but pretreatment with heat shock also significantly shortened recovery time.

We observed that the point at which *X. laevis* tadpoles and 1dpf zebrafish become paralyzed is around 25 MPa (65). This pressure corresponds to approximately the same depth, 2-3km, that some of the deepest diving whales, such as the beaked and sperm whales, can reach (66). This hints at an evolutionary constraint imposed by pressure on surface-dwelling organisms. While activation of heat shock enhanced survival in blastula-staged embryos and aided recovery from paralysis to pressures up to 25 MPa, animals were still paralyzed. This indicates that other evolutionary adaptations are required for surface animals to reach and survive at greater ocean depths.

This is one of the first studies to examine how vertebrates respond to pressure at the organismal level. There have been some studies examining specific aspects of pressure in cell culture, such as the cytoskeleton and its effects on transcription or translation (33, 42). RNA-sequencing experiments were previously performed on adult Zebrafish using much lower pressures for a shorter duration (15 MPa held for 20 min) (67). This dataset was not included in our analysis because, given the low pressure and short exposure time, we do not anticipate that embryos had sufficient time to acclimate to the pressure. Therefore, we ran our own RNA-seq experiment under conditions more similar to those used for *X. laevis* (25 MPa for 2 hours) and found that, as with the other species we examined, pressure induces a heat shock response.

Many RNA-seq datasets from deep-sea organisms, or those exposed to pressure, are available but were excluded for various reasons. First, we focused on metazoans, so we did not include datasets from unicellular eukaryotes, prokaryotes, or fungi. We did not include data sets for other invertebrate metazoans due to difficulty in obtaining genome data, or that these studies compared related species rather than responses to pressure changes (18, 19, 68, 69). Thus, we limited our comparisons to the invertebrate bilaterians *A. japonicus* and *B. lactea.* Though we were unable to process the dataset ourselves, heat shock proteins were also identified as being induced in datasets from the shrimp *Rimicaris* (19).

As described earlier, pressure is predicted to affect many aspects of biology, such as protein misfolding and changes to membrane dynamics. Many of these effects have been shown to activate HSR (70-72). We suspect that protein misfolding may be the primary cause of death, as most of these other effects, such as changes in membrane curvature, should quickly return to normal once the pressure is released; however, the refolding and synthesis of proteins are much slower processes. We propose a model in which activation of the heat shock pathway in response to pressure upregulates proteins that refold or remove misfolded proteins. This helps embryos recover from pressure-induced damage. It is likely that HSR functions with a currently unknown second pathway to increase pressure tolerance. Unlike treatments with heat or CdCl_2_, overexpressing Hsf1 does not change sensitivity to pressure. There are a number of pathways from E3 ubiquitin ligases to cytochrome P450 proteins that are also upregulated in response to pressure that may function in parallel to HSF to increase pressure tolerance.

We observed seasonal changes in tolerance to pressure. The majority of experiments in this study were conducted over the winter, as pressure tolerance for blastula-staged embryos would increase during the summer months. During the summer months, some embryos were often able to tolerate pressures as high as 60 MPa, even though our frog colony and embryos were maintained at similar temperatures, and light was on a 12 hour cycle. To try to control for this, embryos were kept at 14°C whenever they were not being imaged or experimented on. Seasonal variation suggests that non-temperature factors in the care of adult frogs can affect which maternal RNAs or metabolites are deposited into maturing oocytes, leading to changes in tolerance to pressure. For tadpole-staged embryos, this phenomena was less prevalent.

Our study has established parameters for further studies on tolerance of deep-sea pressures. A number of metabolic pathways were identified in genetic studies, but have never been experimentally tested. These include pathways such as trimethylamine N-oxide (TMAO) and polyunsaturated fatty acid synthesis that may be pressure protective beyond the 45 MPa exposures in our work. Though we are currently unable to visualize embryos under pressure, this work identifies the parameters needed to design that equipment for future work. Future application of these methodologies will guide our understanding of how complex animals can survive and thrive under increased hydrostatic pressure and may yield clinical breakthroughs for treating complications that arise in human divers.

## Methods

### Hydrostatic pressure

Pressure chambers were constructed from 9/16” (HM9) tubing capped with an HF9 cap. (Fig. S1) Each chamber holds ∼3.5 mL of fluid, contiguous with the rest of the system, which holds around 70 ml. 10 to 20 embryos were placed into each chamber for experimentation. An analog gauge was used as verification that the digital readout was correct (Grainger: PRO-301L-254W-01). Due to the 1/4 inch NPT connection, this gauge was not compatible with pressures above 60 MPa. Unless otherwise stated, pressures are gradually increased pressure over 30 minutes to indicated pressures, held the pressure for two hours then returned over 15 minutes to atmospheric levels. For experiments that required isolation, such as drug treatments, embryos were placed into 3mm ID flexible PVC tubing capped with PDMS plugs created by using a 4mm biopsy punch. Then, PVC tube with embryos was placed into the chamber. Due to leaching compounds from the PVC tubing, and absorption of substances into the tubing, tubing was routinely boiled in distilled H_2_O for 30 minutes and stored in distilled water until use. No noticeable difference was observed between embryos in PVC tubing and those placed directly in the chamber. For temperature control experiments, we placed the stainless steel pressure chambers in 1 L beakers of water, which were tightly temperature-controlled using either a custom thermoelectric heat pump or a ThermoCube Solid State Chiller circulating chilled water through a copper coil. For pressure experiments, a Syrixus 65x syringe pump (Teledyne) connected to 316ss tubing (highpressure.com) was used. Unless otherwise noted, pressure was increased over 30 minutes to the indicated pressure, held for 2 hours, then released over 15 minutes.

*Aurelia aurita* experiments were carried out at Marine Biological Laboratory. Jellyfish (approximately 5 cm bell diameter) were obtained from the New England Aquarium and kept in a 17 L jellyfish holding tank (Exotic Aquaculture EA-JCT-17) at room temperature until pressure treatments could be performed. All jellyfish were pressure-treated within 24 hours after arrival. For the pressure treatment, jellyfish were removed from the holding tank two at a time, placed in a 1 L pressure vessel (Parr Instruments 4680-1L), and pressurized in steps of 5 MPa every 7.5 minutes, resulting in an overall pressure increase from 0 to 25 MPa over a 30-minute period. The jellyfish were maintained at 25 MPa for 2 hours, then pressure was released in steps of 1.5 MPa every 1.5 minutes until the pressure vessel reached atmospheric pressure. Within 30 minutes after the treatment concluded, jellyfish were frozen individually in 50 ml centrifuge tubes and held at -80°C and dry ice until RNA could be extracted at the University of Pittsburgh.

### RNAseq/qPCR

For RNAseq and qPCR experiments in *X. laevis*, 10 embryos were treated as indicated, then frozen at - 80°C prior to RNA extraction. Thawed embryos were lysed by pipetting up and down with 100µl TX-Lysis solution (10mM HEPES pH 7.4; 150mM NaCl; 2mM EDTA; 0.5% TritonX-100) supplemented with DNAse. The solution was centrifuged at 15,000 RPM for 5 minutes, and the supernatant was moved to a new tube. 500µl Tri reagent (Invitrogen 15596026) was added to the sample and quickly mixed.

For RNA-seq and qPCR experiments in Zebrafish, embryos were treated as indicated and then frozen at -80°C prior to RNA extraction. Embryos were incubated with 1 ml TRI reagent and macerated with a plastic mortar. 200µl Chloroform was added, centrifuged, and the aqueous phase was moved to a new tube containing DNase.

For RNA-seq in *Aurelia aurita*, samples were treated as indicated and then frozen at -80°C prior to RNA extraction. RNA was purified by a modified protocol described in Bouchard, Michaels, and Brown-Harding, 2020 (73). A 50 ml conical tube containing the frozen Moon Jellies was filled with 8M Urea, 5M LiCl. Moon Jellies were placed on a nutator for 5 hours at room temperature to dissolve, then kept at 4°C overnight to aid in RNA precipitaion. Samples were centrifuged at 4,400 RPM at 4°C for 60 minutes. The supernatant was discarded, and the pellet was washed in 75% EtOH, lightly dried, and resuspended in 2 ml TRI reagent. 500 µl was used for further purification.

All RNAseq samples were purified using the Direct-Zol MiniPrep kit (Zymo R2050) per the manufacturer’s instructions, skipping the DNAse steps. Illumina sequencing of the *Xenopus* samples was performed by Novagene, NovaSeq X Plus Series (PE150), while the Zebrafish and *Aurelia* samples were sequenced by Plasmidsaurus.

FastQ files for all organisms were processed ourselves using the Subio64 platform running fastp, Hisat2, and Stringtie functions. *X. laevis* genome V10.1 obtained from Xenbase was used (43, 74). The *B. lactea* genome was obtained from Dr. Chenguang Feng and Dr. Haibin Zhang at the Institute of Deep-Sea Science and Engineering, and the Center for Ecological and Environmental Sciences, Beijing, PRC (20). The *A. japonicus* and Zebrafish genomes were downloaded from NCBI ASM3797524v1 and GRCz12tu (75, 76). The Aurelia aurita genome was downloaded from https://davidadlergold.faculty.ucdavis.edu/jellyfish/ (77). Though this is not the most up-to-date version of the genome, the ABSv1 is not assembled in a way that is compatible with RNA-seq analysis.

For quantitative real-time PCR, 200µl of chloroform was added to the final Tri-reagent and mixed. The sample was centrifuged at 15,000 RPM for 5 minutes, and the clear aqueous fraction was moved to a new tube. 1 ml isopropanol was added to the sample and centrifuged for 10 minutes, and the pellet was washed with 75% ethanol. RNA was resuspended in water, and DNase. The DNase was heat-inactivated at 75°C 15 minutes. cDNA synthesis was performed using the iScript cDNA synthesis kit (Bio-Rad #1708890), and qPCR was run using iTaq SYBR green Supermix (Bio-Rad #1725121) and indicated primers. All RNA-seq experiments are averages of 3 independent experiments conducted on 3 different days.

Hsp70-F GAAGAGTATGCCTTCCAGCAGA

Hsp70-R GGCCTGAGCACCACAGCTAGAAC

Eef1a-F ACCCTCCTCTTGGTCGTTTT

Eef1a-R TTTGGTTTTCGCTGCTTTCT

### Comparative Transcriptomics

To find genes that are conserved across multiple species. We obtained published RNA-seq datasets from NCBI *A. japonicus* (PRJNA507056) (16) and *B. lactea* (PRJNA765439) (17). FastQ files were processed using the same pipeline as the *X. laevis* RNA-seq dataset. Homologous genes were identified by performing a reciprocal BLAST, using the local BLAST software. <https://blast.ncbi.nlm.nih.gov/doc/blast-help/downloadblastdata.html#blast-executables>. Positive gene pairs were defined as any BLAST result with an E-value <0.02.

### Imaging

Confocal images were collected using a Leica SP5 scanning laser confocal or spinning disk confocal with automated microscope software (Micromanager 2.0)(78, 79) controlling an inverted compound microscope (Leica Dmi8) with a 25x/0.95NA water immersion objective lens, equipped with a scanhead (Yokogawa CSUX1-M1L-E) and a CCD camera (Hamamatsu EM-CCD). White light dissection scope images were taken using Micromanager 2.0, acquiring images from a FLIR Blackfly S U3-51S5C, or Blackfly S U3-51S5BW camera on an Olympus SZX7 microscope.

To immobilize tadpoles for time-lapses, a mold was made using a 3D-printed Zebrafish injection mold (https://www.thingiverse.com/thing:4968318) provided by the Dr. Daniel Wagner lab. This mold was used to create PDMS (Electron Microscopy Sciences Sylgard-184 24236-10) channels that held tadpoles in place. For antibody staining, embryos were fixed in 3% TCA for 30 minutes, followed by blocking in CAS Block solution and staining with 1:100 αCytokeratin type II (1H5, DSHB) and 1:1000 αCtnnb1 (C2206, Milipore). GoatαMouse::Alexa-488 and GoatαRabbit::Alexa-647(Invitrogen) were used to detect primary antibodies. For phalloidin staining, embryos were fixed in 3.7% formaldehyde, followed by incubation with Acti-stain 488 phalloidin (Cytoskeleton PHDG1-A). The epidermis was manually removed using fine forceps.

### Movement Analysis and Electrostimulation

Movement was measured from time-lapse sequences using a custom ImageJ macro as described in Davidson et al. (2006) (80). For the exact code, see Figure S6. To stimulate muscle contraction, a Transcutaneous Electrical Nerve Stimulation (TENS) device (TENS 7000, Compass Health Brands Corp.) was connected to a 316 stainless steel wire embedded in a PDMS plate. Using the mold described above to create grooves with a wire exposed at the bottom. Sequential RGB images were collected every 100 ms over 100 seconds with a color camera; only the red channel was used for movement analysis.

For Zebrafish recovery from pressure experiments, the same PDMS-coated plate was used. At least 9 embryos per trial were measured across 3 trials on 3 different days, for a minimum of 27 embryos. Embryos from each trial came from the same clutch and were imaged simultaneously. To calculate the half-time to recovery, we modified the method used for calculating FRAP recovery time. Embryos were grouped by day, and a 15 minute rolling average was plotted. The ½ maximum movement intensity was calculated by taking the average movement index from 500 to 550 seconds, then dividing by 2. A trendline was fitted to the data, and the time at which the trendline crossed the ½ maximum movement intensity was used as our t½.

Zebrafish embryos were heat shocked by placement on a hotplate set to 34°C for 1 hour, then 37°C for 4 hours, and finally 40°C for 30 minutes. Zebrafish were taken directly from the hotplate to the pressure chamber.

### Plasmid Constructs

Hsf1 was PCR-amplified from cDNA generated from RNA isolated from untreated late neurula stage 18 embryos using primers hsf1-F gtgcaggcctctcgagatggacccccacgggacttgtg, and hsf1-R tagttctagaactagtttaggagatgctggagcctgctgg, then cloned into a pCS2 plasmid containing a c-terminal fluorophore. A R79E mutation was introduced by PCR SOEing. pCs2-Etmb::2xmCherry was a gift from Dr. Anne Miller lab (81), krt8::GFP constructs were a gift from Victoria J Allan (37).

### Animals

Adult *X. laevis* were obtained from *Xenopus*1 and maintained under the University of Pittsburgh, Division of Laboratory Animal Resources, under IACUC protocol #24014521 (PHS Assurance Number: D16-00118). Oocytes were collected and fertilized using standard procedures (82, 83). Unless otherwise stated, all embryos were cultured at 14°C.

AB strain zebrafish were obtained from Columbia University as part of a project at the University of Pittsburgh to grow wildtype Zebrafish from around the world. They were maintained under the University of Pittsburgh, Division of Laboratory Animal Resources, under IACUC protocol #24014521 (PHS Assurance Number: D16-00118).

## Terms

HSR: Heat Shock Response pathway
Barotrauma: Physical damage from rapid changes in pressure
TMAO: Trimethylamine N-oxide
Pressure: Hydrostatic pressure.
MZT: maternal-to-zygotic transition

## Acknowledgements

This project was performed with inspiration from the Bio-inspired material architectures for deep sea (BIMADS) group and early advice from Dr. Jeffrey Brodsky. We thank Dr. Jay Green and Dr. Alfredo Garcia for the loan of a pressurizable imaging chamber.

---

**Table 1:** Excel sheet listing the relative expression of genes for the RNAseq experiments performed in this study.

**Movie 1:** Death by pressure occurs by cellular lysis. Left: Control embryos; right: pressure-treated.

**Movie 2:** Death due to pressure in tadpole-stage embryos.

**Movie 3:** Movement analysis of electro-stimulated tadpoles.

**Movie 4:** Heat shock improves recovery after exposure to 25MPa of pressure.

## References

1. T. T. Sutton, R. J. Milligan, "Deep-Sea Ecology" in Encyclopedia of Ecology (Second Edition), B. Fath, Ed. (Elsevier, Oxford, 2019), 10.1016/B978-0-12-409548-9.11010-3, pp. 35-45.

2. M. E. Gerringer, On the Success of the Hadal Snailfishes. Integr Organism Biol 1 (2019).

3. K. Taira, S. Kitagawa, T. Yamashiro, D. Yanagimoto, Deep and Bottom Currents in the Challenger Deep, Mariana Trench, Measured with Super-Deep Current Meters. Journal of Oceanography 60, 919–926 (2004).

4. J. E. Smiley, M. A. Drawbridge, Techniques for live capture of deepwater fishes with special emphasis on the design and application of a low-cost hyperbaric chamber. J Fish Biol 70, 867–878 (2007).

5. 5. N. L. C. McCarthy et al., "Chapter Three - The effect of hydrostatic pressure on lipid membrane lateral structure" in Methods in Enzymology, T. Baumgart, M. Deserno, Eds. (Academic Press, 2024), vol. 700, pp. 49-76.

6. J. R. Winnikoff et al., Homeocurvature adaptation of phospholipids to pressure in deep-sea invertebrates. Science 384, 1482–1488 (2024).

7. M. E. Corkins et al., Similar biomolecular constraints drive convergent adaptation to extreme cold and high pressure. Integr Comp Biol 10.1093/icb/icaf052 (2025).

8. R. C. Dougherty, Temperature and pressure dependence of hydrogen bond strength: A perturbation molecular orbital approach. The Journal of Chemical Physics 109, 7372–7378 (1998).

9. J. Roche et al., Cavities determine the pressure unfolding of proteins. Proc Natl Acad Sci U S A 109, 6945–6950 (2012).

10. C. Charlier et al., Study of protein folding under native conditions by rapidly switching the hydrostatic pressure inside an NMR sample cell. Proc Natl Acad Sci U S A 115, E4169–E4178 (2018).

11. M. L. Sogin et al., Microbial diversity in the deep sea and the underexplored "rare biosphere". Proc Natl Acad Sci U S A 103, 12115–12120 (2006).

12. E. C. Miller et al., Alternating regimes of shallow and deep-sea diversification explain a species-richness paradox in marine fishes. Proc Natl Acad Sci U S A 119, e2123544119 (2022).

13. W. Xu et al., Chromosome-level genome assembly of hadal snailfish reveals mechanisms of deep-sea adaptation in vertebrates. eLife 12, RP87198 (2023).

14. Y. Mu et al., Whole genome sequencing of a snailfish from the Yap Trench (∼7,000 m) clarifies the molecular mechanisms underlying adaptation to the deep sea. PLoS Genet 17, e1009530 (2021).

15. M. E. Gerringer et al., Habitat influences skeletal morphology and density in the snailfishes (family Liparidae). Front Zool 18, 16 (2021).

16. L. Liang, J. Chen, Y. Li, H. Zhang, Insights into high-pressure acclimation: comparative transcriptome analysis of sea cucumber Apostichopus japonicus at different hydrostatic pressure exposures. BMC Genomics 21, 68 (2020).

17. G. Yan et al., Comparative transcriptomic analysis of in situ and onboard fixed deep-sea limpets reveals sample preparation-related differences. iScience 25, 104092 (2022).

18. J. Chen, H. Liu, S. Cai, H. Zhang, Comparative transcriptome analysis of Eogammarus possjeticus at different hydrostatic pressure and temperature exposures. Sci Rep 9, 3456 (2019).

19. J. Zhang, Q. L. Sun, Z. D. Luan, C. Lian, L. Sun, Comparative transcriptome analysis of Rimicaris sp. reveals novel molecular features associated with survival in deep-sea hydrothermal vent. Sci Rep 7, 2000 (2017).

20. R. Liu et al., De Novo Genome Assembly of Limpet Bathyacmaea lactea (Gastropoda: Pectinodontidae): The First Reference Genome of a Deep-Sea Gastropod Endemic to Cold Seeps. Genome Biol Evol 12, 905–910 (2020).

21. E. M. Case, J. B. Haldane, Human physiology under high pressure: I. Effects of Nitrogen, Carbon Dioxide, and Cold. J Hyg (Lond) 41, 225–249 (1941).

22. R. W. Brauer, Hydrostatic pressure effects on the central nervous system: perspectives and outlook. Philos Trans R Soc Lond B Biol Sci 304, 17–30 (1984).

23. A. G. Macdonald, I. Gilchrist, K. T. Wann, A. E. Wilcock, "THE TOLERANCE OF ANIMALS TO PRESSURE" in Animals and Environmental Fitness: Physiological and Biochemical Aspects of Adaptation and Ecology, R. Gilles, Ed. (Pergamon, 1980), 10.1016/B978-0-08-024938-4.50027-6, pp. 385-403.

24. K. K. Jain, High-pressure neurological syndrome (HPNS). Acta Neurologica Scandinavica 90, 45–50 (1994).

25. S. E. Wilcock, K. T. Wann, A. G. Macdonald, The motor activity of Crangon crangon subjected to high hydrostatic pressure. Marine Biology 45, 1–7 (1978).

26. K. K. Jain, High-pressure neurological syndrome (HPNS). Acta Neurol Scand 90, 45–50 (1994).

27. N. Zahn et al., Normal Table of Xenopus development: a new graphical resource. Development 149 (2022).

28. P. D. Nieuwkoop, J. Farber, Normal Table of Xenopus Laevis (Daudin) (North-Holland Publishing Company, American Elsevier Publishing Co., 1975).

29. 29. J. Yang, T. Aguero, M. L. King, "Chapter Eight - The Xenopus Maternal-to-Zygotic Transition from the Perspective of the Germline" in Current Topics in Developmental Biology, H. D. Lipshitz, Ed. (Academic Press, 2015), vol. 113, pp. 271-303.

30. R. Keller, D. Shook, Dynamic determinations: patterning the cell behaviours that close the amphibian blastopore. Philos Trans R Soc Lond B Biol Sci 363, 1317–1332 (2008).

31. J. Holtfreter, Die totale Exogastrulation, eine Selbstablösung des Ektoderms vom Entomesoderm. *Wilhelm Roux’* Archiv für Entwicklungsmechanik der Organismen 129, 669–793 (1933).

32. T. Elsdale, D. Davidson, Timekeeping by frog embryos, in normal development and after heat shock. Development 99, 41–49 (1987).

33. B. Bourns, S. Franklin, L. Cassimeris, E. D. Salmon, High hydrostatic pressure effects in vivo: Changes in cell morphology, microtubule assembly, and actin organization. Cell Motility 10, 380–390 (1988).

34. H. C. Crenshaw, J. A. Allen, V. Skeen, A. Harris, E. D. Salmon, Hydrostatic pressure has different effects on the assembly of tubulin, actin, myosin II, vinculin, talin, vimentin, and cytokeratin in mammalian tissue cells. Exp Cell Res 227, 285–297 (1996).

35. H. W. Detrich, 3rd, S. K. Parker, R. C. Williams, Jr., E. Nogales, K. H. Downing, Cold adaptation of microtubule assembly and dynamics. Structural interpretation of primary sequence changes present in the alpha- and beta-tubulins of Antarctic fishes. J Biol Chem 275, 37038–37047 (2000).

36. G. Li, J. K. Moore, Microtubule dynamics at low temperature: evidence that tubulin recycling limits assembly. Mol Biol Cell 31, 1154–1166 (2020).

37. E. J. Clarke, V. J. Allan, Cytokeratin intermediate filament organisation and dynamics in the vegetal cortex of living Xenopus laevis oocytes and eggs. Cell Motil Cytoskeleton 56, 13–26 (2003).

38. K. Faire et al., E-MAP-115 (ensconsin) associates dynamically with microtubules in vivo and is not a physiological modulator of microtubule dynamics. J Cell Sci 112 (Pt 23), 4243–4255 (1999).

39. J. Riedl et al., Lifeact: a versatile marker to visualize F-actin. Nat Methods 5, 605–607 (2008).

40. B. Peters, J. Kirfel, H. Bussow, M. Vidal, T. M. Magin, Complete cytolysis and neonatal lethality in keratin 5 knockout mice reveal its fundamental role in skin integrity and in epidermolysis bullosa simplex. Mol Biol Cell 12, 1775–1789 (2001).

41. P. Vijayaraj, G. Sohl, T. M. Magin, Keratin transgenic and knockout mice: functional analysis and validation of disease-causing mutations. Methods Mol Biol 360, 203–251 (2007).

42. 42. A. M. Zimmerman, "High-Pressure Studies in Cell Biology" in International Review of Cytology, G. H. Bourne, J. F. Danielli, K. W. Jeon, Eds. (Academic Press, 1971), vol. 30, pp. 1-47.

43. A. M. Session et al., Genome evolution in the allotetraploid frog Xenopus laevis. Nature 538, 336–343 (2016).

44. B. B. Mughal, M. Leemans, P. Spirhanzlova, B. Demeneix, J. B. Fini, Reference gene identification and validation for quantitative real-time PCR studies in developing Xenopus laevis. Sci Rep 8, 496 (2018).

45. T. Ishikawa, T. Igarashi, K. Hata, T. Fujita, c-fos induction by heat, arsenite, and cadmium is mediated by a heat shock element in its promoter. Biochem Biophys Res Commun 254, 566–571 (1999).

46. Y. Morishima et al., Regulation of cytochrome P450 2E1 by heat shock protein 90-dependent stabilization and CHIP-dependent proteasomal degradation. Biochemistry 44, 16333–16340 (2005).

47. V. P. Androutsopoulos, A. M. Tsatsakis, D. A. Spandidos, Cytochrome P450 CYP1A1: wider roles in cancer progression and prevention. BMC Cancer 9, 187 (2009).

48. W. Sang, W. H. Ma, L. Qiu, Z. H. Zhu, C. L. Lei, The involvement of heat shock protein and cytochrome P450 genes in response to UV-A exposure in the beetle Tribolium castaneum. J Insect Physiol 58, 830–836 (2012).

49. S. Khan, J. J. Heikkila, Distinct patterns of HSP30 and HSP70 degradation in Xenopus laevis A6 cells recovering from thermal stress. Comp Biochem Physiol A Mol Integr Physiol 168, 1–10 (2014).

50. J. T. Young, J. J. Heikkila, Proteasome inhibition induces hsp30 and hsp70 gene expression as well as the acquisition of thermotolerance in Xenopus laevis A6 cells. Cell Stress Chaperones 15, 323–334 (2010).

51. A. Ali, S. Bharadwaj, R. O’Carroll, N. Ovsenek, HSP90 interacts with and regulates the activity of heat shock factor 1 in Xenopus oocytes. Mol Cell Biol 18, 4949–4960 (1998).

52. J. J. Heikkila, N. Ohan, Y. Tam, A. Ali, Heat shock protein gene expression during Xenopus development. Cell Mol Life Sci 53, 114–121 (1997).

53. T. Elsdale, M. Pearson, M. Whitehead, Abnormalities in somite segmentation following heat shock to Xenopus embryos. J Embryol Exp Morphol 35, 625–635 (1976).

54. J. P. Woolfson, J. J. Heikkila, Examination of cadmium-induced expression of the small heat shock protein gene, hsp30, in Xenopus laevis A6 kidney epithelial cells. Comp Biochem Physiol A Mol Integr Physiol 152, 91-99 (2009).

55. S. Gordon, S. Bharadwaj, A. Hnatov, A. Ali, N. Ovsenek, Distinct stress-inducible and developmentally regulated heat shock transcription factors in Xenopus oocytes. Dev Biol 181, 47–63 (1997).

56. M. Bienz, Developmental control of the heat shock response in Xenopus. Proceedings of the National Academy of Sciences 81, 3138–3142 (1984).

57. S. D. Westerheide, J. Anckar, S. M. Stevens, Jr., L. Sistonen, R. I. Morimoto, Stress-inducible regulation of heat shock factor 1 by the deacetylase SIRT1. Science 323, 1063–1066 (2009).

58. G. Herbomel et al., Dynamics of the full length and mutated heat shock factor 1 in human cells. PLoS One 8, e67566 (2013).

59. C. Chaudhary, A. J. Richardson, D. S. Schoeman, M. J. Costello, Global warming is causing a more pronounced dip in marine species richness around the equator. Proc Natl Acad Sci U S A 118 (2021).

60. R. S. Brennan et al., Experimental evolution reveals the synergistic genomic mechanisms of adaptation to ocean warming and acidification in a marine copepod. Proc Natl Acad Sci U S A 119, e2201521119 (2022).

61. P. A. Cleves, C. J. Krediet, E. M. Lehnert, M. Onishi, J. R. Pringle, Insights into coral bleaching under heat stress from analysis of gene expression in a sea anemone model system. Proc Natl Acad Sci U S A 117, 28906–28917 (2020).

62. K. Kaarniranta et al., Hsp70 accumulation in chondrocytic cells exposed to high continuous hydrostatic pressure coincides with mRNA stabilization rather than transcriptional activation. Proc Natl Acad Sci U S A 95, 2319–2324 (1998).

63. K. Kaarniranta et al., Stress responses of mammalian cells to high hydrostatic pressure. Biorheology 40, 87–92 (2003).

64. D. Cottin et al., Sustained hydrostatic pressure tolerance of the shallow water shrimp Palaemonetes varians at different temperatures: insights into the colonisation of the deep sea. Comp Biochem Physiol A Mol Integr Physiol 162, 357–363 (2012).

65. M. P. Ford, S. P. Colin, J. H. Costello, Effects of high hydrostatic pressure on the mechanical performance and behavior of shallow-water jellyfish (Aurelia aurita). J Exp Biol 229 (2026).

66. E. E. Henderson, S. W. Martin, R. Manzano-Roth, B. M. Matsuyama, Occurrence and Habitat Use of Foraging Blainville’s Beaked Whales () on a US Navy Range in Hawaii. Aquat Mamm 42, 549–562 (2016).

67. M. L. Hu et al., Gene expression responses in zebrafish to short-term high-hydrostatic pressure. Zool Res 43, 188–191 (2022).

68. J. W. Chen, H. L. Liu, S. Y. Cai, H. B. Zhang, Comparative transcriptome analysis of at different hydrostatic pressure and temperature exposures. Sci Rep-Uk 9 (2019).

69. Z. Gan et al., Comparative transcriptomic analysis of deep- and shallow-water barnacle species (Cirripedia, Poecilasmatidae) provides insights into deep-sea adaptation of sessile crustaceans. BMC Genomics 21, 240 (2020).

70. G. Balogh et al., Heat stress causes spatially-distinct membrane re-modelling in K562 leukemia cells. PLoS One 6, e21182 (2011).

71. Z. Bromberg, Y. Weiss, The Role of the Membrane-Initiated Heat Shock Response in Cancer. Front Mol Biosci 3, 12 (2016).

72. B. Zhang, Y. Fan, K. Tan, HSF1 Activation Mechanisms, Disease Roles, and Small Molecule Therapeutics. Int J Biol Sci 21, 3351–3378 (2025).

73. C. Bouchard, J. Michaels, H. Brown-Harding, RNA isolation from corals and other cnidarian species using urea-LiCl as a denaturant. Anal Biochem 588, 113472 (2020).

74. S. Chu et al., Xenbase: 25 years of integrating molecular and biomedical data from Xenopus. Genetics 232 (2026).

75. L. Sun, C. Jiang, F. Su, W. Cui, H. Yang, Chromosome-level genome assembly of the sea cucumber Apostichopus japonicus. Sci Data 10, 454 (2023).

76. R. E. Broughton, J. E. Milam, B. A. Roe, The complete sequence of the zebrafish (Danio rerio) mitochondrial genome and evolutionary patterns in vertebrate mitochondrial DNA. Genome Res 11, 1958–1967 (2001).

77. D. A. Gold et al., The genome of the jellyfish Aurelia and the evolution of animal complexity. Nat Ecol Evol 3, 96–104 (2019).

78. A. D. Edelstein et al., Advanced methods of microscope control using muManager software. J Biol Methods 1 (2014).

79. A. Edelstein, N. Amodaj, K. Hoover, R. Vale, N. Stuurman, Computer control of microscopes using microManager. Curr Protoc Mol Biol **Chapter 14**, Unit14 20 (2010).

80. L. A. Davidson, M. Marsden, R. Keller, D. W. Desimone, Integrin alpha5beta1 and fibronectin regulate polarized cell protrusions required for Xenopus convergence and extension. Curr Biol 16, 833–844 (2006).

81. G. von Dassow, K. J. Verbrugghe, A. L. Miller, J. R. Sider, W. M. Bement, Action at a distance during cytokinesis. J Cell Biol 187, 831–845 (2009).

82. H. Sieve, Xenopus: A Laboratory Manual (Cold Spring Harbor Press, 2023).

83. H. L. Sive, R. M. Grainger, R. M. Harland, Early Development of Xenopus laevis: A Laboratory Manual (2000).

